# Direct and indirect cues can enable dual-adaptation, but through different learning processes

**DOI:** 10.1101/2021.04.09.439164

**Authors:** Marion Forano, Raphael Schween, Jordan A. Taylor, Mathias Hegele, David W. Franklin

## Abstract

Switching between motor tasks requires accurate adjustments for changes in dynamics (grasping a cup) or sensorimotor transformations (moving a computer mouse). Dual-adaptation studies have investigated how learning of context-dependent dynamics or transformations is enabled by sensory cues. However, certain cues, such as color, have shown mixed results. We propose that these mixed results may arise from two major classes of cues: “direct” cues, which are part of the dynamic state and “indirect” cues, which are not. We hypothesized that explicit strategies would primarily account for adaptation for an indirect color cue but would be limited to simple tasks while a direct visual separation cue would allow implicit adaptation regardless of task complexity. To test this idea, we investigated the relative contribution of implicit and explicit learning in relation to contextual cue type (colored or visually shifted workspace) and task complexity (one or eight targets) in a dual-adaptation task. We found that the visual workspace location cue enabled adaptation across conditions primarily through implicit adaptation. In contrast, we found that the color cue was largely ineffective for dual-adaptation, except in a small subset of participants who appeared to use explicit strategies. Our study suggests that the previously inconclusive role of color cues in dual-adaptation may be explained by differential contribution of explicit strategies across conditions.

**New & Noteworthy:** We present evidence that learning of context-dependent dynamics proceeds via different processes depending on the type of sensory cue used to signal the context. Visual workspace location enabled learning different dynamics implicitly, presumably because it directly enters the dynamic state estimate. In contrast, a color cue was only successful where learners were apparently able to leverage explicit strategies to account for changed dynamics. This suggests a unification for the previously inconclusive role of color cues.

## Introduction

Humans perform a variety of motor tasks, switching effortlessly between them as they go through their daily lives. Switching between these tasks requires accurate adjustments for changes in task dynamics caused by object properties (grasping a cup) or sensorimotor transformations (moving a computer mouse). These adjustments are associated with the formation and selection of appropriate motor memories (1). In order to investigate the formation and recall of these independent motor memories, dual-adaptation studies expose participants to opposing dynamics (2–9) or opposing visuomotor transformations (10–20).

Dual-adaptation studies have found that simultaneous adaptation is contingent on the presence of sensory cues that allow the sensorimotor system to identify the context-dependent changes in dynamics (21–30). Within the range of these contextual cues that have been examined, some are clearly effective, such as different workspace locations (visual or physical) (28, 31) or distinct lead-in or follow-through movements (25, 32, 33). On the other hand, other contextual cues have either shown less effectiveness or provided mixed results. For example, some studies have found color cues to effectively enable dual-adaptation (7, 34–36), while others have not (3, 31). The reason behind these mixed findings remains unclear.

A priori, there is no reason to expect that the ability to use different cues to associate task dynamics with specific contexts should differ depending on the type of cue, as long as the cues are similarly salient and predictive of the dynamics. To explain why color cues could be treated differently than for example lead-in movements, it has been suggested that motor adaptation is only sensitive to cues that pertain to the dynamic state of the body or manipulated object (31, 37–39) and are therefore, part of a relevant neural preparatory state (40, 41). Under this view, cues like lead-in movements, workspace location, arm posture (3, 42, 43) or the used effector constitute “direct” or “primary” cues, containing direct information about the dynamic state of the body. Other cues like color (37), or sequence order (4) are “indirect” or “conditional” representations. These are not immediately implicated in the dynamic state and therefore an association to context-dependent dynamics needs to be formed through a different process.

Here, we hypothesize that the effectiveness of direct versus indirect cues may map onto implicit and explicit motor learning processes (44–46), which have gained recent attention in visuomotor adaptation studies (47, 48). Indeed, recent work suggests that dual-adaptation to opposing cursor rotations can arise from implicit motor adaptation as well as explicit cognitive strategies, with implicit dual-adaptation enabled only by cues that can be considered immediately relevant to the dynamic state of the body, such as the use of separate hands or movement directions (11–13, 16, 49). Given that explicit cognitive strategies have been suggested to contribute to force field learning under some circumstances (28, 50–53), differential contributions of explicit and implicit learning under specific task conditions could then potentially explain the conflicting findings regarding the effectiveness of color cues outlined above. One candidate for task conditions that could influence the contribution of explicit strategies is the complexity of the task. By this broad term, we refer to factors inherent in the task that require potential solution strategies to be more or less elaborate. For example, we would consider a task with one target direction to have lower complexity than a task with several different target directions. This is because a single target direction allows counteracting a force field by a simple heuristic, such as to deliberately “push left” in order to counter a clockwise curl field. This strategy would work with a single, distal target but fail if the task contains other targets, in which case a “push counterclockwise” strategy would be required. Such a counterclockwise strategy would be more elaborate (assuming that participants plan in cartesian coordinates (54), in that it requires extracting information from multiple targets in its generation and adapting the direction of pushing to the specific target in its application. A more complex task may therefore put more strain on cognitive resources like working memory and reduce the contribution of capacity-limited explicit learning. This interpretation might explain the mixed findings of the effectiveness of a color cue across different tasks.

Therefore, here we investigate the relative contribution of explicit and implicit components to the adaptation of opposing force fields using a two-by-two design of contextual cue type and task complexity. For cue type, we compared a direct state representation (lateral visual shift of the workspace) with an indirect cue (different background colors). For task complexity, we compared a simple task (reaching in one direction) with a complex task (reaching in eight directions). We hypothesized that a direct state representation would lead to automatic or implicit dual-adaptation while indirect cues would not enable it. Moreover, explicit strategies would be expected to contribute most within a simple task.

## Materials and Methods

### Participants

A total of 30 (19 male, 11 female, mean age 27 ± 0.8 years) force field naive participants took part in two experiments (six participants in each of 5 conditions, described below). All participants were right-handed based on the Edinburgh handedness questionnaire (55), and provided written informed consent before participation. The Ethics Committee of the Medical Faculty of the Technical University of Munich approved the study.

### Apparatus

Seated in a custom adjustable chair, participants made right-handed reaching movements while grasping the endpoint of a vBOT planar robotic manipulandum. Their forearm was supported against gravity using an air sled. The vBOT system is a custom built robotic interface (56) that can apply state-dependent forces on the handle while recording the position and velocity as participants move in a planar workspace located approximately 15 cm below the shoulder (Fig 1C). A six-axis force transducer (ATI Nano 25; ATI Industrial Automation) measured the end-point forces applied by the participant on the handle. Joint position sensors (58SA; Industrial encoders design) on the motor axes were used to calculate the position of the vBOT handle. Position and force data were sampled at 1kHz. Visual feedback to the participants was provided horizontally from a computer monitor (Apple 30” Cinema HD Display, Apple Computer, Cupertino, CA, USA; response time: 16 ms; resolution: 2560 × 1600) fixed above the plane of movement and reflected via a mirror system that prevented visual feedback of the participants’ arm. The visual feedback, provided in the same plane as the movement, consisted of circles indicating the start, target and cursor positions on a black background. Participants wore headphones to remove environmental noise and provide necessary auditory information.

**Figure 1.**
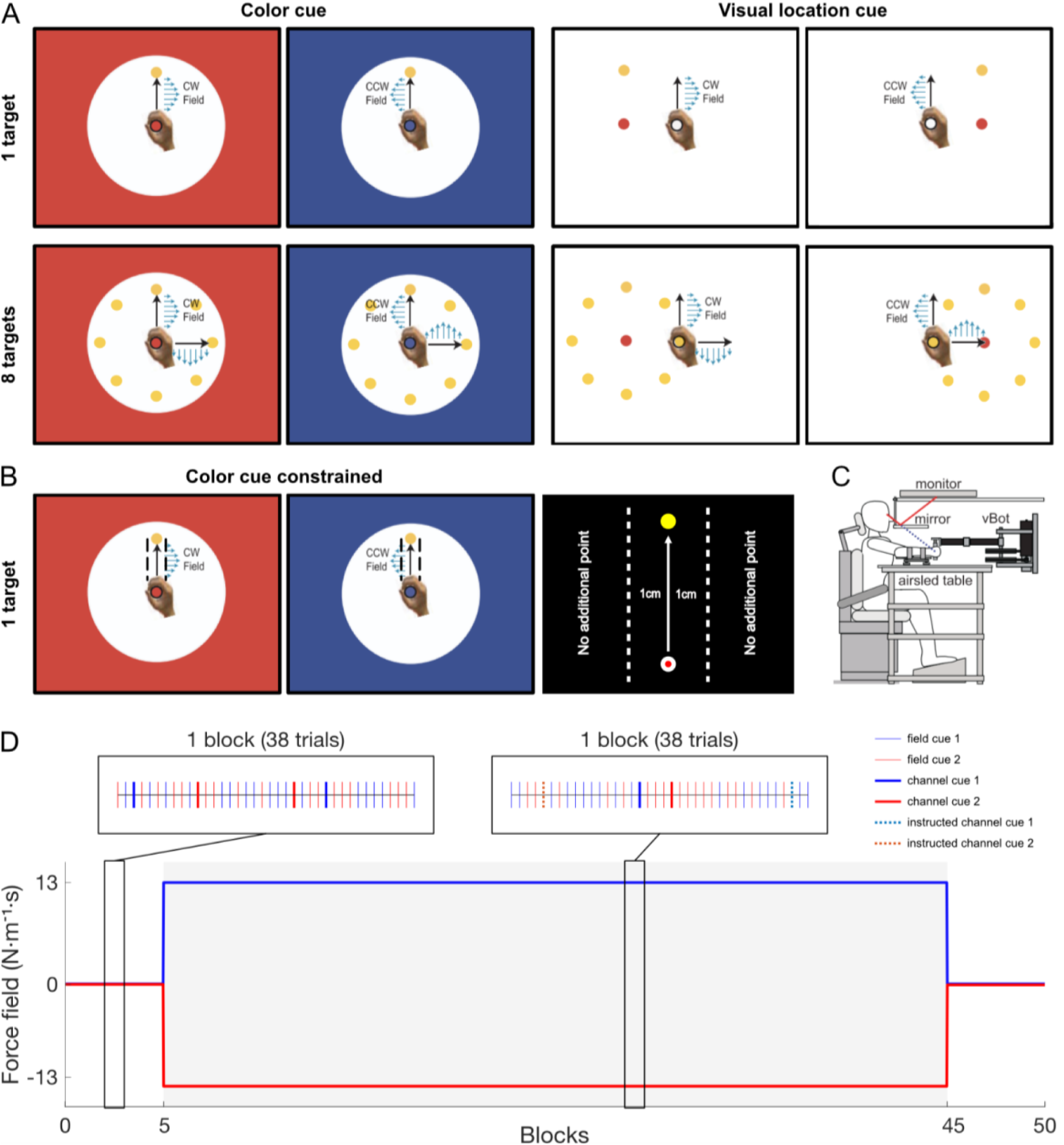
Experimental Setup. **A**. Workspace layouts of the four conditions of experiment 1: Color-1 (top left), Location-1 (top right), Color-8 (bottom left) and Location-8 (bottom right). Participants always physically performed reaching movements in the center of the workspace. During these physical movements, the visual feedback was presented either in red and blue for the color cue experiments or left and right (±10cm) for the visual location cue experiments. Each of these two cues were associated with either a clockwise (CW) or counter clockwise curl (CCW) force field. In the 8 targets experiments, two examples of reaching movements (forward, 0° and right, 90°) are displayed. In the 8 targets experiments, only one target appeared on each trial. **B**. Layout of the experiment 2, Color-1-constrained, with the kinematic constraints displayed (dotted black lines, which were not shown to the participant). If the participant’s hand crossed a virtual channel of 1cm on each side of the target trajectory, no increase in score was considered. **C**. Participants were seated and grasped the handle of the robotic manipulandum (vBOT) with their forearm supported by an airsled. Visual feedback of movements, displayed by the monitor, were viewed through a mirror so that they appear in the plane of movement. **D**. Temporal paradigm of the experiment, including the structure of pseudo-randomized blocks. The grey area represents the exposure phase.

### Protocol

Participants initiated a trial by moving the cursor representing the hand position (red circle of 1.0 cm diameter) into the start position (grey circle of 1.5 cm diameter) located in the middle of the screen, approximately 20 cm directly in front of the participant. This start position turned from grey to white once the cursor entered it. Once the hand was within the start position for a random time exponentially distributed between 1 and 2s, the target (yellow circle of 1.5 cm diameter) appeared, prompting participants to initiate a reaching movement. The target was located 10.0 cm from the start position (the exact location of the target depended on the condition). The movement was considered complete when the participants maintained the cursor within the target for 600ms in a maximum time of 2s after target appearance. After each trial, the participant’s hand was passively driven to the start position while visual feedback regarding the success of the previous trial was provided. Successful trials were defined as trials where participants hit the target without overshooting and where their movement’s peak speed was comprised between 37 and 53 cm/s. On these trials, the participants received positive feedback (e.g., “great” for peak speeds between 41 and 49 cm/s or “good” for peak speeds between 37 and 41 or 49 and 53 cm/s), and a counter displayed on the screen increased by one point (score). In contrast, messages of “too fast” or “too slow” were provided when the peak speed exceeded 53 cm/s or did not reach 37 cm/s, respectively. “Overshoot” feedback was provided when the cursor overshot the target by more than 1 cm. Finally, a low tone was played on unsuccessful trials. Movements were self-paced, as participants were able to take a break before starting the next trial. Short breaks were enforced every 115 trials throughout each session, except for the first break which was delayed to 153 trials to narrow the time span to the exposure phase. Participants were instructed to reach naturally straight to the target right after its appearance (go-cue) on each trial. During each movement, the vBOT was either passive (null field), produced a clockwise (CW) or counterclockwise (CCW) velocity-dependent curl force field, or produced a mechanical channel. For the velocity-dependent curl field (3), the force at the handle was given by:

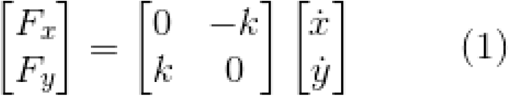

where k was set to either ±13 N·m^− 1^·s, with the sign of k determining the force field direction (CW or CCW). On a channel trial, the reaching movement was confined to a simulated mechanical channel with a stiffness of 6000 N·m^− 1^ and damping of 2 N·m^− 1^·s acting lateral to the line from the start to the target (57, 58). These “channel” trials were used to measure the lateral force, produced by the participants on the wall of this mechanical channel, that reflects their compensation to the exposed force field.

### Instructions

In addition to ordinary channel trials, some channel trials, called “instructed” trials, were combined with a clear auditory instruction. Participants were informed that the environmental disturbance was removed for a trial when they heard the message ‘Robot off’. This audio message was provided before the start of the instructed trial, during the passive return portion of the movement, giving the participant enough time to process it and act accordingly. These instructed trials played a key role in the experiment as they allowed us to assess explicit strategies employed by participants (53). Therefore, to make sure that the participants understood and followed the instructed trials properly, written instructions were given just before the start of the experiment. In addition, to further remind them of the instruction, these instructions were provided a second time during the last break of the pre-exposure phase (one block before switching to the exposure phase) and a third time on the first break following the passage into the exposure phase (two blocks after switching phases). The same written instructions were provided either in English or German, depending on the preference of the subject and varied slightly depending on the phase of the experiment. Finally, a questionnaire was provided after the experiment in order to assess participants’ awareness of the task and their explicit knowledge about any strategies they may have employed to counteract the force fields. The purpose of the questionnaire was to have a descriptive comparison with participants’ performance, and to help us classify them as ‘‘aware’’. Participants who reported recognizing a pattern of two opposing, different, or left and right directions of force under a given cue either correctly or with reversed association (i.e. still identifying that there were a “left” and “right” field and that these corresponded to the two cues) were defined as being “aware”.

### Experimental Paradigm

#### Experiment 1

In this study, participants were required to complete a simple dual-adaptation paradigm using a two-by-two design of contextual cue type and task complexity (Fig 1). Participants were presented with two opposing force fields, a CW or CCW curl force field coupled with specific contextual cues, to investigate their adaptation. Two different types of cues were used: an indirect cue and a direct cue. The indirect cue, used in conditions Color-1 and Color-8, was set as an environmental color cue. In these conditions, the workspace (background and cursor) was colored in either blue or red, with the color determining the field direction (Fig 1A, left side, top and bottom). With a colored background, a black circle (15cm diameter) was displayed in the middle of the screen to allow participants to clearly see the start position, cursor and target. The direct cue, used in conditions Location-1 and Location-8, was a visually shifted workspace. In these conditions, movements were always made in the same central hand position, but the entire visual workspace (center starting location, targets and cursor) was shifted to the left (−10cm from the sagittal axis) or to the right (+10cm from the sagittal axis) of the midline, with the offset’s direction determining the field direction (Fig 1A, right side, top and bottom). The cues were displayed throughout the experiment. For task complexity conditions, one forward target (10cm to the front) was used in conditions Color-1 and Location-1, whereas eight different target directions (separated by 45 degrees) were used in conditions Color-8 and Location-8.

#### Experiment 2

An additional experiment was designed to confirm the effect of the explicit component in learning a dual-adaptation task. This experiment was identical to condition Color-1 except for a supplementary constraint on the perpendicular kinematic error. The error range was set at ±1cm from the direct straight line to the target such that if the cursor position in the x-direction crossed the limit while reaching to the target, the trial was set as unsuccessful, no specific feedback and no increase in score was observed (Fig 1B). Additional feedback was provided depending on the overall performance (speed, overshoot). Participants were informed, in the instruction preceding the experiment, that the absence of feedback meant that they were not straight enough and had to correct their trajectory to gather points.

Each experiment started with 5 blocks in the pre-exposure phase (190 trials) where the same null force field was associated with the cue. In the written instructions, this phase was referred to the participants as the “Robot Off” phase. This was followed by an exposure phase consisting of 40 blocks (1520 trials) where one contextual cue (e.g. +10cm visual shift or blue workspace) was associated with one force field (e.g. CW), whereas the other contextual cue (e.g. -10 cm visual shift or red workspace) was associated with the other force field (e.g. CCW, Fig 1). Each condition was counterbalanced such that half of the participants experienced this phase with contextual cues matched to one set of force field directions, whereas the other half of the participants had contextual cues matched to the opposing force field directions. Finally, a post-exposure phase of 5 blocks (190 trials) was completed where, again, a null force was applied suddenly, without further instruction.

The experimental blocks consisted of 38 trials each, evenly split between the two contextual cues. Out of these 19 trials for each cue, 17 movements were performed in either a null field (pre-exposure phase), CW or CCW force field (exposure phase). In the one target conditions, these trials were always performed in the forward direction, whereas in the eight targets conditions, movements were performed to each of the eight targets: three trials in the forward direction and two trials in all other directions. Two trials (out of 19) were a channel trial and an instructed channel trial, which were always performed in the forward direction. Within each block, trials were pseudo randomized ensuring that the channel trials and instructed channel trials never directly followed one another. Overall, the five conditions were identical except for the number of target directions (one or eight), the cue type (colored workspace or shifted workspace), and the spatial constraint introduced in experiment 2.

### Analysis

Data were analyzed offline using MATLAB (2018b, The MathWorks, Natick, MA). All measurements were low-pass filtered at 40 Hz using a 10^th^ order zero-phase-lag Butterworth filter (filtfilt). Over the course of the experiment, the lateral force measured can vary due to the natural drift of the mass of the arm over the airsled. In order to avoid interference in our measurements from this drift, we subtracted the mean offset in lateral force for each participant measured between 250 and 150 ms prior to the movement start, from the total force trace. This is of particular importance in these experiments as participants could potentially be attempting to reduce errors through an initial force bias which could then influence measures of force compensation. The start of the reaching movement was defined as the cursor leaving the start position (cursor center crossing the start position’s 1.5cm-radius) and the end was defined as the cursor entering the target (cursor center crossing the target’s 1.5cm-radius). To measure adaptation and its origin (implicit or explicit strategy), we examined the following variables: kinematic error, force compensation, relative lateral force and reaction time. Here we focus on three different stages of the experiment: pre-exposure (all 5 blocks), late exposure (last 5 blocks) and post-exposure (all 5 blocks).

#### Kinematic error

To measure the kinematic error, we calculated the maximum perpendicular error (MPE) on each null or curl field trial. MPE is defined as the signed maximum perpendicular distance between the hand trajectory and the straight line joining the start and the end of the current displayed target. For completeness, we also calculated the kinematic error at the time of peak velocity and at a fixed time early in the movement (200 ms after movement start).

#### Force compensation

To measure adaptation, we calculated the force compensation by regressing the end-point force applied by the participant on the handle (lateral measured force) against the force needed to compensate perfectly for the force field (59). Specifically, the slope of the linear regression through zero is used as a measure of force compensation. The perfect compensatory force was determined as the forward velocity of the current trial multiplied by the force field constant *k*. As each condition was counterbalanced across participants the values were flipped such that the force compensation associated with each cue has the same sign for all participants. Since the measure of force compensation is relative to the field, we inverted the sign of cue 2’s data for plotting in order to visually differentiate the two opposing forces.

#### Relative lateral force

To examine the shape and timing of the force applied by the participant to compensate for the disturbance, the relative lateral forces were calculated. Individual force trials were aligned to peak velocity, and clipped between -300 and +300 ms from this peak. For averaging and visualization, we normalized forces in the x-axis by the peak of the perfect compensation force profile. This perfect force profile was calculated as the y-axis velocity multiplied by the force field constant *k*.

#### Reaction time

As an additional indicator of explicit strategies in the adaptation to the opposing force-fields, the reaction time was calculated for each trial and compared between trial types (field, channel and instructed trials) across experiment phases. Reaction time was calculated as the time between the go-cue (target appearance) and the start of the movement (cursor leaving the target). For a fair comparison across the conditions, we examine only the reaction times in the forward reaching directions.

### Statistics

We performed statistical analysis using JASP (version 0.13.1). For each condition, we calculated the mean of force compensation across the two cues (cue 1 and cue 2) for the final level of adaptation (mean across 5 last blocks of exposure phase) and post-exposure phase (mean across all 5 blocks). Except for statistics on the pre-exposure phase, the mean value of force compensation across pre-exposure phase (mean across all 5 blocks) was subtracted.

To assess adaptation in each condition, these baseline-corrected force compensation values were compared against zero using one sample t-tests (in the case of normally distributed data) or Wilcoxon tests (when not normally distributed). In order to investigate the implicit and explicit contributions, paired t-tests were performed to assess differences in force compensation between channel and instructed channel trials. Additional Wilcoxon tests were performed for force compensation on instructed channel trials for final level of adaptation only. Finally, we performed a two-way ANOVA to compare the effects of *Cue type* (Color or Visual location) and *Complexity* (1 target or 8 targets) on force compensation (channel trials only) for the final level of adaptation only.

We quantified the reaction times for the three types of trials (field trial, channel trial and instructed channel trial) over the exposure period to look for differences (Fig 7F) using the linear mixed models approach with main effects of conditions (5 levels) and trial type (3 levels), and Participant as a random effect.

## Results

In this study, we investigated the mechanisms involved in dual-adaptation to force-fields. Specifically, we built an experimental design to assess the degree of implicit and explicit processes in adaptation, and varied the complexity of the task and type of the contextual cues to understand the relevance of these contributions in different conditions. In experiment 1, four conditions tested the four possible combinations: Color cue one target (Color-1; indirect, low complexity), Color cue eight targets (Color-8; indirect, high complexity), Visual workspace location cue one target (Location-1; direct, low complexity) and Visual workspace location cue eight targets (Location-8; direct, high complexity). In each of these conditions, participants performed reaching movements towards one or eight targets in an identical pre-exposure, exposure and post-exposure phase.

### Adaptation

In each condition, kinematic errors (Fig 2A-D, e.g. Supplemental Fig. S1 (https://doi.org/10.6084/m9.figshare.16566225)) and force compensation (Fig 2E-H) remained close to zero in the pre-exposure phase (one sample t-test, for force compensation: Color-1: t_5_=0.289, p=0.784; Color-8: t_5_=0.765, p=0.479; Location-1: t_5_=-1.349, p=0.235; Location-8: t_5_=-0.177, p=0.867; and for kinematic errors: Color-1: t_5_=-0.085, p=0.935; Color-8: t_5_=2.019, p=0.099; Location-1: t_5_=60.823, p=0.448; Location-8: t_5_=-0.027, p=0.980), with no clear differences between the two cues. We observed an increase in kinematic error from pre-exposure to exposure phase, demonstrating that participants did not expect the introduction of the force fields. However, only visual location conditions showed a strong decrease of kinematic error from early to late exposure phase (Fig 2B,D) and a strong increase of force compensation on channel trials (Fig 2F,H). A Wilcoxon test revealed a significant increase in Location-1 (V=21.000, p=0.031) up to 80±6% (mean ± s.e.m.) and in Location-8 (V=21.000, p=0.031, here and below, identical test statistics are reflective of the limited number of rankings available with Wilcoxon test and our sample size) up to 60±8%. These results demonstrate the presence of dual-adaptation for participants in visual location conditions. In contrast, both color conditions do not display a clear decrease in kinematic error (Fig 2A,C) or increase in force compensation (Fig 2E,G) over the exposure phase. Indeed, no significant increase in force compensation was found for Color-1 (V=18.000, p=0.156), with participants producing up to 24±16% adaptation, and for Color-8 (V=3.000, p=0.156), with -4±2% force compensation in final adaptation. These results suggest that participants in color conditions were not able to adapt to the two opposing force fields. However, it is important to note the high variability across participants on both the kinematic error and force compensation for Color-1 condition. This variability and the final level of adaptation of 24±16% provide questions about whether only certain individuals show adaptation. We find similar results for kinematic errors at the time of peak velocity (e.g. Supplemental Fig. S2 (https://doi.org/10.6084/m9.figshare.16566225) and at 200 ms into the movement (e.g. Supplemental Fig. S3 (https://doi.org/10.6084/m9.figshare.16566225), as well as for force compensation without subtracting the bias force prior to movement start (e.g. Supplemental Fig. S4 (https://doi.org/10.6084/m9.figshare.16566225).

**Figure 2.**
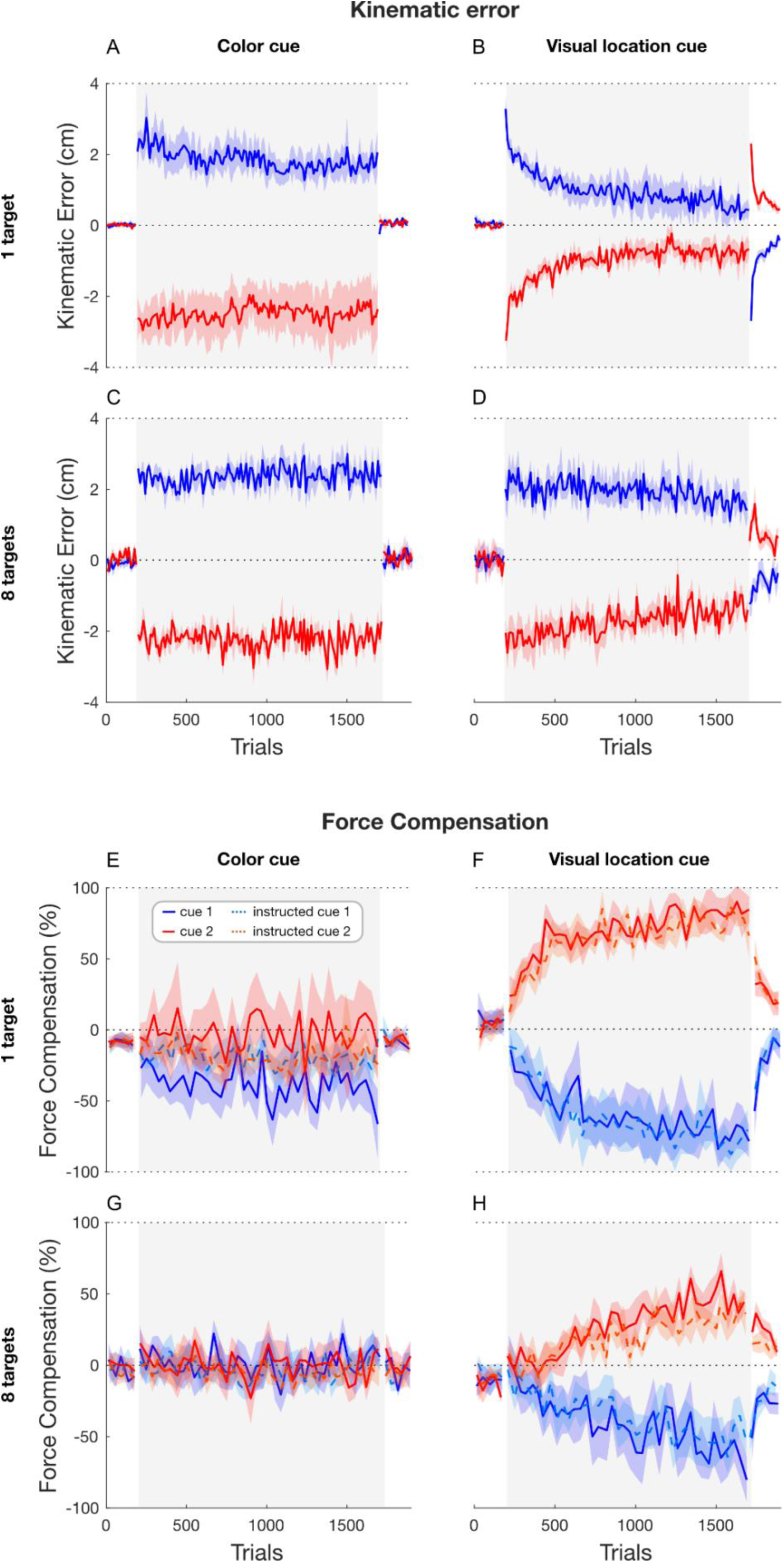
Overall adaptation for experiment 1. The mean (solid line) and s.e.m. (shaded region) across participants is represented for cue 1 (blue color) and cue 2 (red color) for the pre-exposure (190 trials), exposure (grey area, 1520 trials) and post-exposure (190 trials) phases. The adaptation is represented by the kinematic error (**A-D)**, measured as Maximum Perpendicular Error (MPE) and the force compensation (**E-H)** for the four conditions. Normal channel trials (red and blue solid lines) and instructed channel trials (light red and blue dotted lines) are both presented.

### Comparison across conditions

In order to test the effects of cue type and task complexity across all four conditions, we ran a two-way ANOVA to compare the effects of *Cue type* (Color or Visual location) and *Complexity* (1 target or 8 targets) on force compensation (channel trials only). Comparing the final level of adaptation (5 last blocks of exposure phase), we first found a significant effect for cue type (F_1,20_=40.048, p< .001). These results show that the visual workspace location is a more effective cue than the colored workspace for dissociating force fields. Second, a significant effect was found for complexity (F_1,20_=9.466, p=0.006). That is, decreasing the number of target directions increased the amount of adaptation over the same number of training trials. There was no significant interaction (F_1,20_=0.048, p=0.829).

### Implicit and explicit contributions

In order to differentiate implicit and explicit contributions to the adaptation of these force fields, we performed Wilcoxon tests on instructed channel trials for the final level of adaptation (mean across cue 1 and cue 2). Force compensation on instructed channel trials revealed a significant adaptation for both Location-1 (V=21.000, p=0.031) and Location-8 conditions (V=21.000, p=0.031). Furthermore, the adaptation on instructed channel trials was similar to those on the channel trials for both visual location conditions. For Location-1, participants produced a force compensation of 78±6% in instructed channel trials (Fig 2F) compared to 80±6% in channel trials (no significant difference with paired Wilcoxon test: W=6, p=0.438). Similarly, in Location-8, participants produced 45±7% force compensation in the instructed channel trials (Fig 2H) compared to 60±8% in channel trials (no significant difference with paired test: t_5_=-2.094, p=0.090). That is, participants produced similar levels of force compensation in both visual workspace location conditions on the instructed channels despite being notified that the force field was not present on these trials. These results suggest that adaptation to the opposing force fields was likely implicit. In contrast, force compensation on instructed channel trials revealed no significant adaptation for both Color-1 (V=12.000, p=0.844) and Color-8 conditions (V=7.000, p=0.563). Even though the force compensation level on channel trials reached 24±16% in Color-1 condition, the force compensation on the instructed channel trials stayed close to zero (1±3%, one sample t-test: t_5_=0.465, p=0.661), suggesting that, if any adaptation was present, this adaptation was likely explicit. As expected, as no adaptation was present in the Color-8 condition (−4±2%, one sample t-test: t_5_=-2.129, p=0.086), no force compensation was seen on the instructed channel trials (−3±3%, one sample t-test: t_5_=-0.959, p=0.382).

Additionally, participants demonstrated clear after-effects in both visual workspace location conditions, with an increase in kinematic error in the opposite direction (0.8±0.05cm for one target, 0.6±0.1cm for eight targets) and a force compensation significantly different from zero (Wilcoxon test, Location-1: V=21.000, p=0.031; Location-8: V=21.000, p=0.031), both of which persisted over the entire 5 blocks of post-exposure. In contrast, participants in both color cue conditions showed no after-effects, with no significant difference from zero (Wilcoxon test, Color-1: V=11.000, p=1.000; Color-8: V=14.000, p=0.563), suggesting no overall adaptation. Again, this highlights the effectiveness of visual workspace location for dual-adaptation and, together with the previous results, suggests that this may arise through implicit adaptation.

### Individual results

The mean participants’ results across the four conditions show clear differences across task complexity and cue type. However, there is a large variability in force compensation for the Color-1 target experiment, making it difficult to investigate the force compensation on the instructed trials and the implication of implicit or explicit adaptation. Consequently, we examined the individual force compensation profiles (Fig 3) in order to distinguish between the impacts of the explicit and implicit components. In the visual workspace location cue conditions, all of the participants that showed independent adaptation to each force field based on visual inspection (10 out of 12) also showed no clear difference in force compensation between the channel and instructed channel trials. In contrast, 11 out of 12 participants in the Color Cue conditions (all 6 in the eight target and 5 in the one target) showed no independent adaptation to the two force fields. The remaining participant (P1) was able to adapt correctly to the force fields in the Color-1 condition. Interestingly, P1 adapted almost immediately and showed complete adaptation on channel trials while the force compensation of instructed channel trials remained close to zero. This contrasts with the participants presented visual workspace location cues, who adapted gradually and showed no clear difference between the channel and instructed channel trials. These results suggest that visual workspace location cue’s adaptation may be mainly implicit whereas P1’s adaptation under color cues, may be explicit.

**Figure 3.**
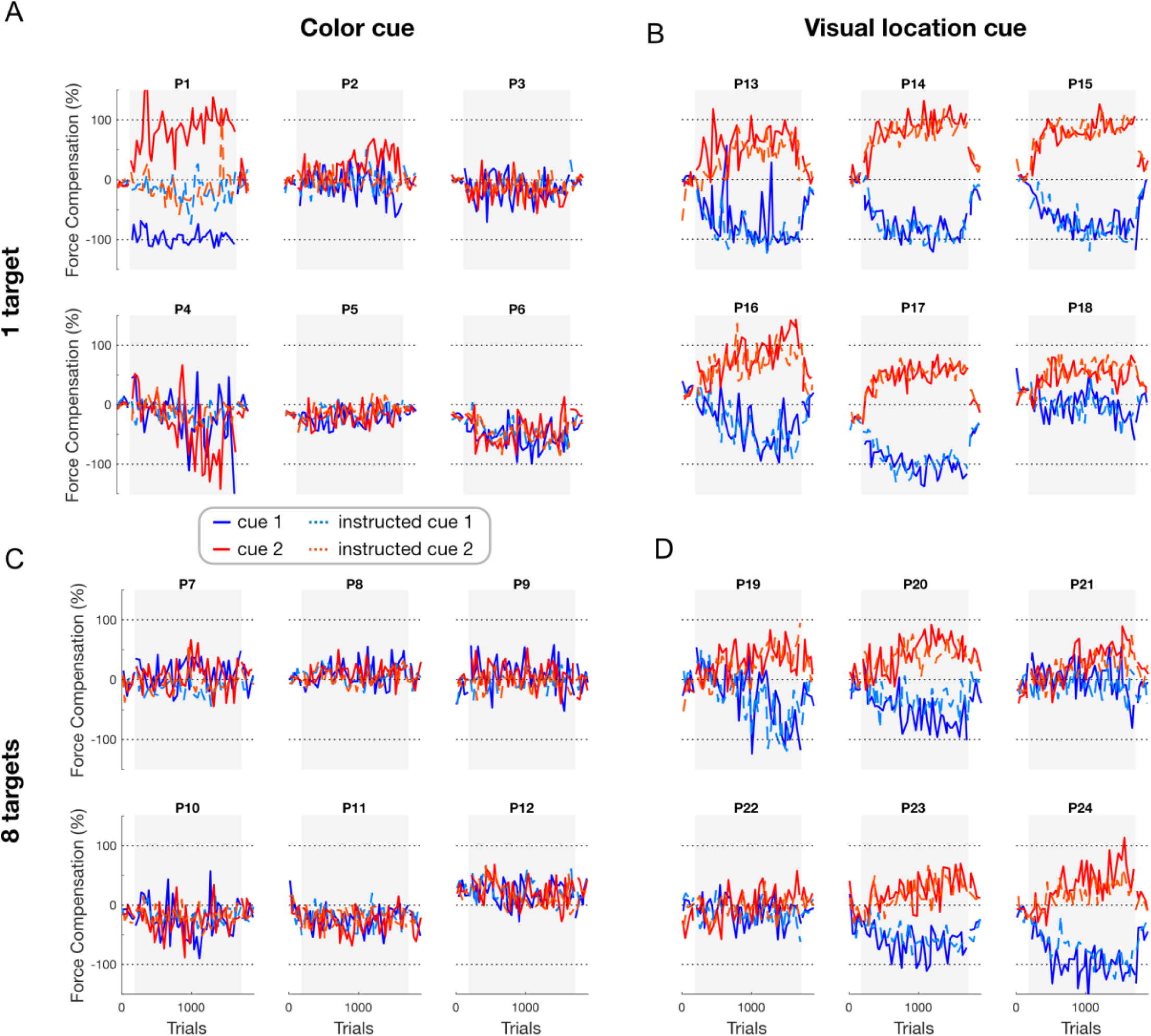
Individual force compensation on the normal channel trials (red and blue solid lines) and the instructed channel trials (light red and blue dotted lines) for Color-1 (**A**), Location-1 (**B**), Color-8 (**C**), and Location-8 conditions (**D**). The grey area represents the exposure phase.

One of the key results of experiment 1 is that only one out of the six participants in the Color-1 condition displayed evidence of dual-adaptation. Moreover this participant showed evidence of a different approach than participants from visual workspace location cue conditions, suggesting explicit adaptation (P1). The fact that only one participant adapted is surprising, as several previous studies have suggested that color cues can be used for dual-adaptation when moving in a single direction (7, 34, 35). A possibility is that other factors besides complexity modulate learning in this condition. Our hypothesis that dual-adaptation with color cues is mediated by explicit strategies provides a possible explanation: whereas implicit learning proceeds regardless of any intention, whether or not learners express explicit strategies likely depends on their perceived necessity to do so (60). Our standard force field task can in principle be solved largely by feedback-based corrections, since reward depends on meeting the velocity criterion and eventually arriving at the target, but not on anticipatory compensation of the force field. We introduced an additional experiment with a single condition (Color cue constrained one target) to test for this possibility. Reasoning that emphasizing the reduction of movement curvature would encourage participants to utilize an anticipatory strategy, we introduced a spatial accuracy constraint: the score was only increased when the participant’s movements were within 1cm of the straight line joining the start and target positions. We predicted that this manipulation would increase explicit learning.

### Experiment 2

Experiment 2 was similar to the Color-1 condition of experiment 1, using color cues to detect opposing force fields in reaching a single forward target. However, an additional horizontal kinematic constraint was imposed to emphasize reduction of spatial errors. Similar to previous conditions, kinematic errors (Fig 4A) and force compensation (Fig 4B) remained close to zero in the pre-exposure phase (one sample t-test: t_5_=0.214, p=0.839 for force compensation and t_5_=-0.305, p=0.773 for kinematic error), with no clear differences between the two cues. After an expected sudden increase in kinematic error at the beginning of the exposure phase, we observed a small reduction in kinematic error and a slow increase in force compensation over the exposure phase up to 39±20% (Fig 4B, solid lines). However, a Wilcoxon test revealed no significant increase in force compensation (V=0.123, p=0.907) for the end of exposure phase, possibly due to the high variability between participants. Moreover, we found no evidence of adaptation on instructed channel trials with a mean of 3±6% of force compensation (Fig 4B, dotted lines), confirmed by a Wilcoxon test (V=11, p=1).

**Figure 4.**
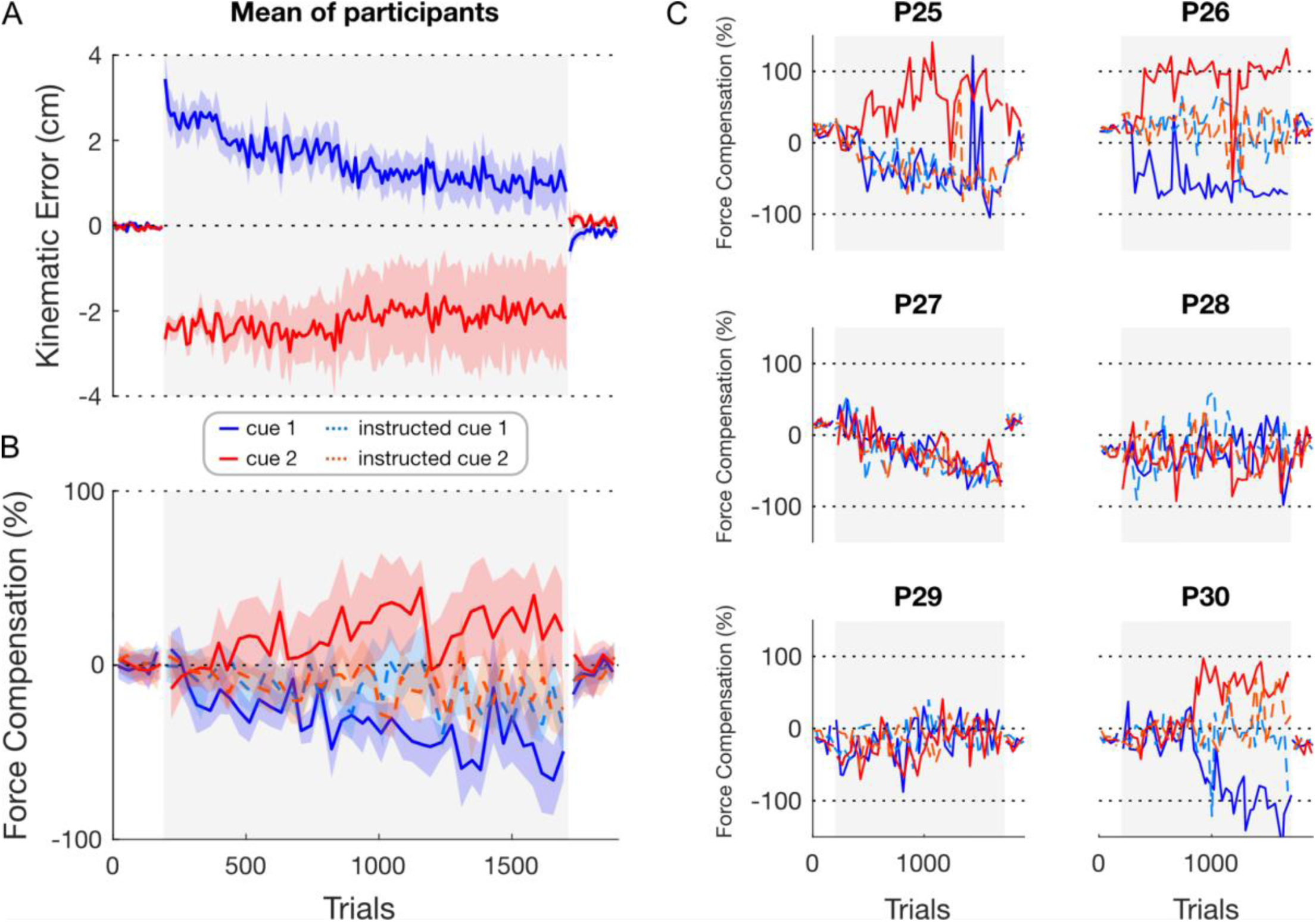
Adaptation for experiment 2. **A**. Kinematic error represented as the mean (solid line) and s.e.m. (shaded region) across participants. **B**. Mean (solid and dotted line) and s.e.m. (shaded region) across participants for force compensation on both the normal (red and blue solid lines) and instructed channel trials (light red and blue dotted lines). **C**. Individual force compensation profiles for each participant. The grey area represents the exposure phase.

Looking at the individual force compensation, three (P25, P26, P30) out of the six participants adapted simultaneously to the opposing force fields (Fig 4C). Similar to participant P1 (Color-1 condition), two of the three participants (P26, P30) showed a rapid change in the predictive force during the exposure phase, as we might expect for an explicit strategy during adaptation. Again, we investigated the presence of force compensation in instructed channel trials to assess the weight of implicit and explicit components during adaptation. P26 and P30 showed a difference between these two types of trials (similar to P1), with almost a full adaptation from channel trials but no observable difference in the predictive force between the two instructed channels, which remained close to zero. We also found no observable difference in the two instructed channel trials for P25, although in this case the force compensation was similar to that of the cue 2 channel trial. The finding that all adapted participants in the color cue conditions showed no clear difference in instructed channels between the two cues suggests that any participants who adapted successfully relied on explicit strategies in this condition.

Additionally, after-effects were absent when the instructed channel trials remained close to zero, but the participants showed adaptation (explicit learners). In contrast, among those participants who adapted implicitly after-effects were consistently present. This is consistent with the notion that after-effects that persist in the presence of errors mainly capture an implicit component of adaptation, which participants cannot simply remove by abandoning an explicit aiming strategy.

### Force Profiles

To further examine adaptation and compare channel and instructed channel trials, we analyzed the force produced by the participants relative to a perfect compensation of the force field applied by the vBot. The force profiles (Fig 5) show similar results as the force compensation (Fig 3, 4), with visual workspace location cue conditions showing adaptation of both channel and instructed channel trials, and no adaptation for the Color cue eight targets condition (Fig 5B). These results are supported by the individual force profiles (Fig 6B, C, D). In the Color Cue one target conditions (Color-1 and Color-1-constrained), we see a small adaptation on the channel trials, whereas the instructed channel trials remain close to zero (Fig 5A, E). As expected, there is a large variability between participants on these two conditions (Fig 6A, E), with participants P1, P25, P26 and P30 showing differences between the channel and instructed channel force profiles, suggesting that an explicit component can contribute to the adaptation when presented with Color Cues.

**Figure 5.**
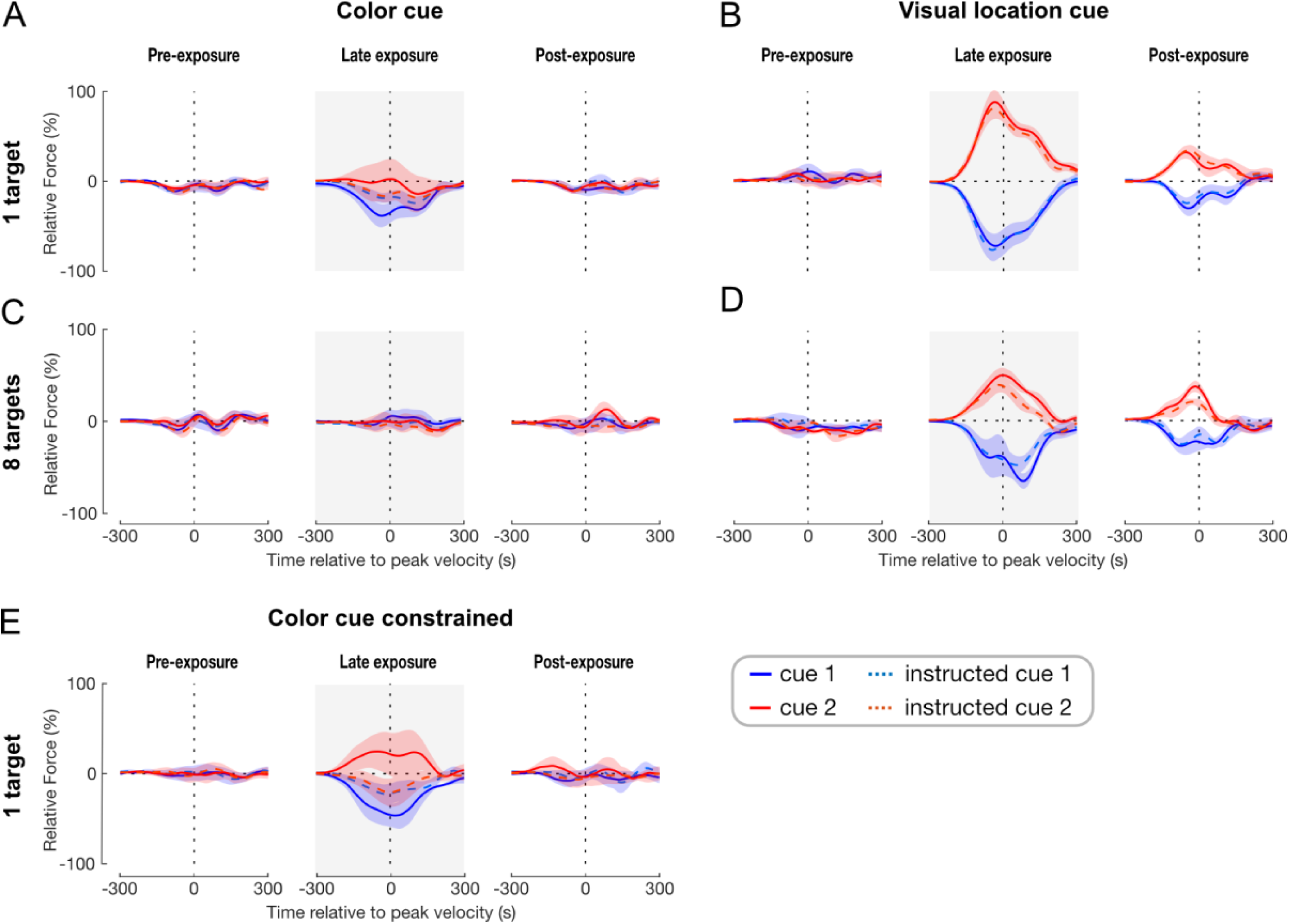
Force profiles as a function of movement time. Channel trials are displayed in red and blue solid lines, and instructed channel trials are displayed in light red and blue dotted lines. The force values are shown as a percentage of perfect force compensation and aligned to peak velocity. For each cue, the mean between participants (solid line) and s.e.m. (shaded region) of force compensation is taken from all trials in each phase (pre-exposure (all 5 blocks), exposure (grey area, final 10 blocks) and post-exposure (all five blocks) for Color-1 (**A**), Location-1 (**B**), Color-8 (**C**), Location-8 conditions (**D**), and Color-1-constrained (**E**).

**Figure 6.**
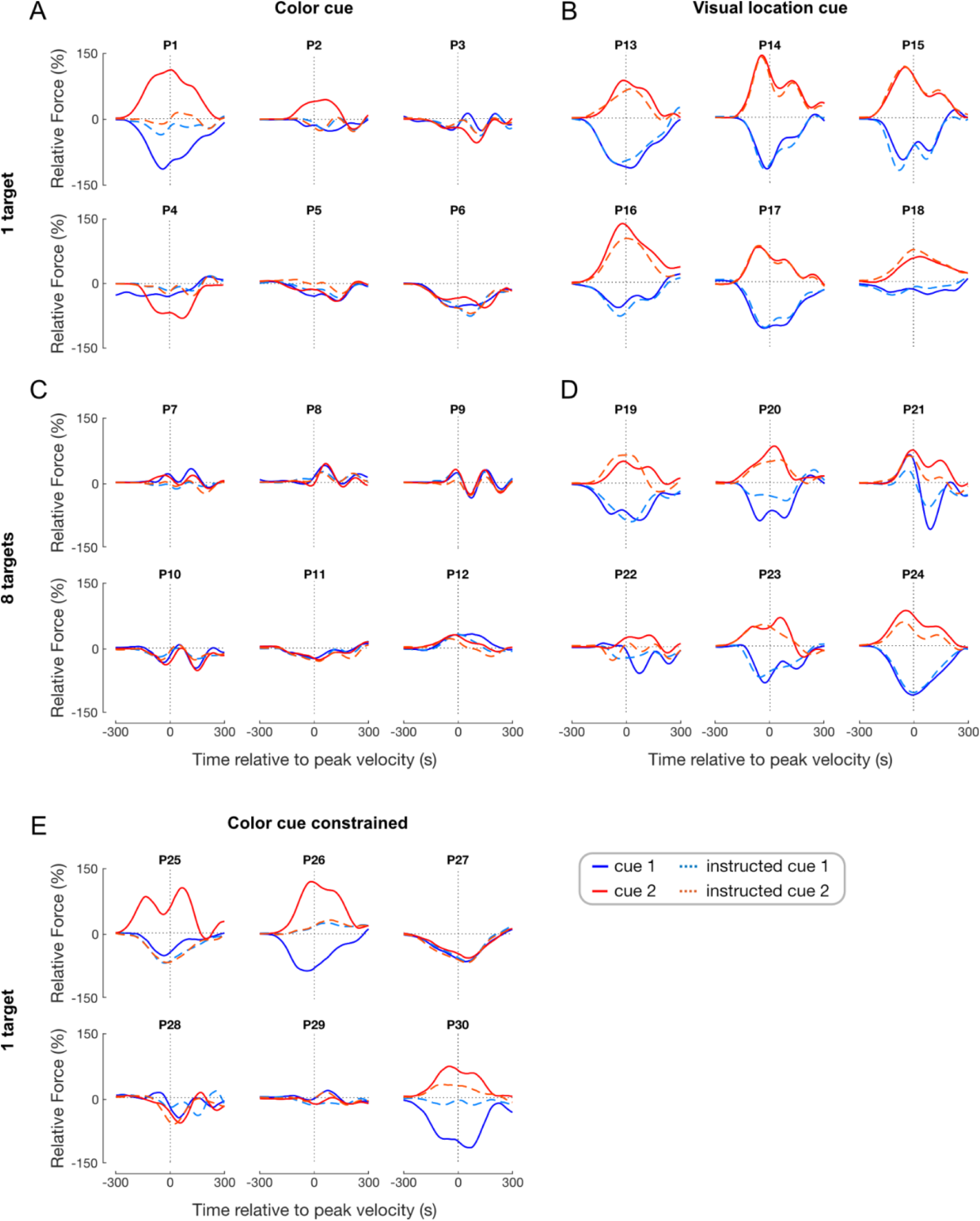
Individual force profiles as a function of movement time for the late exposure phase. Channel trials are displayed in red and blue plain lines, and instructed channel trials in light red and blue dotted lines. For each cue, the mean of force compensation is taken from the final 10 blocks of the late exposure phase for Color-1 (**A**), Location-1 (**B**), Color-8 (**C**), Location-8 conditions (**D**), and Color-1-constrained (**E**).

**Figure 7.**
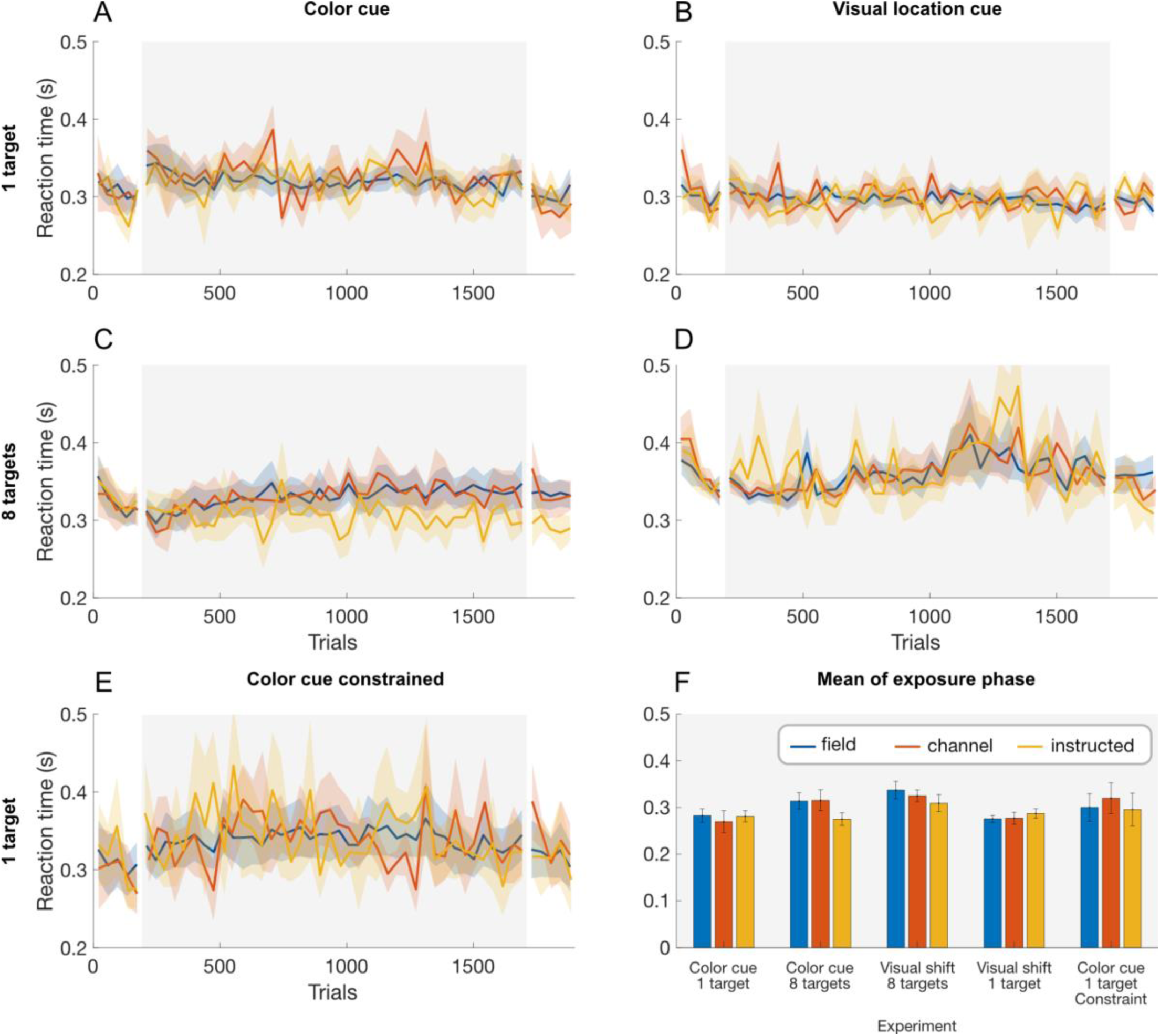
Reaction time according to trial type: field trials (blue), channel trials (orange) and instructed channel trials (yellow). For each trial type, we calculated the mean across participants (solid line) and s.e.m. (shaded region) for Color-1 (**A**), Location-1 (**B**), Color-8 (**C**), Location-8 conditions (**D**), and Color-1-constrained (**E**). **F**. To compare across experiments, the mean and s.e.m are shown across the exposure phase (all 40 blocks). The grey area represents the exposure phase.

### Reaction Time

Reaction time has been shown to be increased with explicit learning strategies (61–64). Therefore, we calculated the reaction time on field trials (null or force field), channel trials and instructed channel trials for forward reaching movements throughout the experiment (Fig 7 A-E). All three trial types showed similar reaction times across the conditions, although there were variations in these values over the time course of the experiment. We quantified the reaction times for these three trial types over the exposure period to look for differences (Fig 7F) using a linear mixed models approach with fixed effects of Condition (5 levels) and Trial type (3 levels), and Participant as a random effect. We found no significant effect of trial type (F_2,50_=0.051, p=0.950), conditions (F_4,25_=0.386, p=0.817) and no trial type*conditions interaction (F_8,50_=2.096, p=0.054). These results are consistent with the finding that the vast majority of participants did not seem to show any indication of used strategy. Interestingly the variability was highest for the Color Cue constrained condition (Fig 7E), suggesting, such as previous results, that a few participants may have used explicit strategies.

### Awareness Assessment

Participants completed a questionnaire from which we assess their awareness of force field directions, use of strategies in the task and self-evaluation of their performance. In the answers reported, we saw a pattern between participants’ performance in the experiment and their awareness of the association between cue and force field direction. We classified participants as “aware” if they reported the direction of the force field under a given cue either correctly or with reversed association (i.e. still identifying that there were a “left” and “right” field and that these corresponded to the two cues)”. Overall, participants P1, P2, P4, P5, P6, P9, P13, P25, P26, P30 were classified as being “aware” of the force directions and their association to a specific cue. In visual location conditions (Location-1 and Location-8), only one participant (P13) was classified as aware, even though most showed clear dual-adaptation. This supports our interpretation that the participants in the visual location conditions primarily adapted implicitly. In color conditions (Color-1, Color-8 and Color-1-constrained), all participants that showed adaptation were also able to report differences in the direction of the force field. Again, this supports our previous interpretation regarding color cues; this indirect cue does not allow implicit adaptation unless participants can leverage explicit knowledge about their environment.

## Discussion

In this study, we investigated how cue type and task complexity influence the contributions of implicit or explicit processes in dual-adaptation to opposing force fields. Five different conditions were set up across two experiments: Color cue one target (indirect, low complexity), Color cue eight targets (indirect, high complexity), Visual workspace location cue one target (direct, low complexity), Visual workspace location cue eight targets (direct, high complexity) and Color cue one target constrained (indirect, low complexity, emphasis on spatial accuracy). Summarizing the overall picture, in experiment 1 we found stronger overall adaptation with the visual workspace location cue than with the color cue, supporting different effectiveness for these cue types as in previous studies (31), and importantly learning was absent in the majority of the participants in the color cue conditions. Less adaptation was also found with eight targets compared to a single target. The absence of a significant interaction gave us no reason to assume that this effect differs between cue types. We acknowledge that the limited sample size per group is a limitation of our study and the latter, group-specific results should be interpreted with particular caution. However, despite individual variations in the amount of adaptation, we found clear, consistent results across all experiments. That is, if any adaptation took place in the color cue experiments, there was evidence that it occurred through explicit adaptation, whereas any adaptation in the visual workspace experiment was associated with evidence for implicit adaptation.

In both experiments, we also examined the contribution of implicit and explicit adaptation to dual-adaptation using instructed channels, where we cued the absence of the force field (53). In the visual workspace location conditions, we found similar levels of force compensation in both the channel trials and instructed channel trials suggesting that this learning arose primarily implicitly. For color cues, individual results showed clear adaptation for four (out of eighteen) participants. However, while these participants displayed force compensation on channel trials, their force compensation on instructed channel trials remained close to zero, suggesting the contribution of explicit strategies.

We now consider how the two different cues in our study (color and visual workspace location) may relate to implicit and explicit learning processes. Contextual cues can vary from direct representations of the body state (28, 42) or object properties (22, 38, 39, 65) to more indirect representations (66–68) such as color (37), or sequence order (4). In the first case, cues are directly related to the current state of the body or properties of the manipulated object and therefore closely representative of the forward dynamics, while in the second case, the relation between the environmental cue and the current state is more conditional. While we believe that both “direct” or “primary” and “indirect” or “conditional” cues can deliver relevant information for dissociating two opposing environments and enable dual-adaptation, we hypothesized that they would do so via different processes, explaining diverging findings regarding their effectiveness (31).

In the visual workspace separation conditions we found clear dual-adaptation across both the one target and eight targets conditions, both on a group and individual level. Significant dual-adaptation on instructed channel trials and persistent after-effects revealed a clear contribution of implicit learning under the visual workspace location cue. This agrees with our premise that visual workspace location is in the class of cues that are directly representative of the dynamic state of the system.

In contrast, we found no evidence of implicit learning in the color cue conditions. Over all three color cue conditions (Color-1, Color-8 and Color-1-constrained), there was little overall adaptation, but this varied considerably between participants. Four out of the eighteen participants exhibited clear dual-adaptation. However, in all four, the difference in adaptation between the instructed and non-instructed channel trials and absence of after effects indicated the use of explicit strategies. Furthermore, all four participants showed awareness that the cue color was associated with different disturbances, even though no information was provided prior to the experiment, which may be a further indication that learning in these participants was explicit. These results provide support for our hypothesis: that color cues are ineffective in eliciting implicit dual-adaptation, at least within the duration of our experiment.

What role does task complexity play for these implicit and explicit routes of dual-adaptation? A key characteristic of explicit processing is that it is capacity-limited (45, 69). We reasoned that such capacity limitations would lead to learners having a higher probability of finding a useful explicit strategy in simple tasks where straightforward heuristics, such as pushing in one direction, would yield immediate success on each trial. This probability would be lower on complex tasks that require a more elaborate rule, such as pushing counterclockwise. Such an expectation is in line with findings that explicit learning is faster with fewer targets in visuomotor learning (70). Although our current results show an effect of task complexity across cue types, this appears to be carried largely by a single individual, rendering the results on task complexity inconclusive. Our findings thus do not lend strong support to the hypothesis that task simplicity alone would compel learners to use explicit strategies. Reasoning that emphasizing spatial accuracy might focus the attention of the participants on reducing kinematic errors and boost explicit strategies (71) we added experiment 2. This manipulation, emphasizing spatial accuracy, appeared to increase explicit learning, with three out of six participants showing evidence of explicit strategies. Overall, these results provide support for the hypothesis that dual-adaptation with color cues is mediated by explicit learning depending on task conditions, with no evidence of implicit dual-adaptation in any of the 18 participants in the color cue conditions.

Previous findings provided mixed results on the effectiveness of color cues for dual-adaptation with some studies showing adaptation whereas others finding no effect. Among those studies that found color cues effective in enabling dual-adaptation to force fields, the majority tested task conditions such as reaching towards a single target or in a single-joint movement (7, 34, 35) that presumably facilitate the use of explicit strategies by affording simple, heuristic solutions (for example pushing to the left or right of the reaching movement). In contrast, studies finding no dual-adaptation contained multiple targets and full-arm movements (3, 31). A notable exception from this pattern is a study by Osu and colleagues (36). In their study, two cues (visual and audio) were displayed in addition to the colored background: a windmill-like diagram illustrating the force field that would be applied and a tone signalling the direction of the force field. Here, several factors may have facilitated explicit adaptation, including the visual presentation of the force field and long inter-trial intervals allowing ample planning of compensatory movements (64, 72). The study furthermore showed that dual-adaptation depended on randomized presentation of the contexts. Whereas increased randomness might be suspected to conflict with explicit strategy use (73), we note that the presence of color cues in the study by Osu and colleagues would still have allowed successful strategy application. Furthermore, subsequent studies have not found randomized presentation to be sufficient in eliciting dual-adaptation with color cues (31), suggesting that other factors may play a role. Overall, our work shows that color cues do not enable learning via the same process as more direct cues (including visual workspace location), and may, at least in early learning stages, rely on explicit strategies.

Notably, our reasoning assigns different mechanisms to a potential effect of adapting under low versus high complexity (one or eight targets) with explicit and implicit learning, and by extension under the visual workspace location or color cue. For implicit learning (visual workspace location cue), we assume that reduced adaptation with more targets results from effectively reducing the number of practice trials per target. Limited spatial generalization (74– 76) then predicts that adaptation will be slower with more targets. In contrast, for explicit learning (mainly found with the color cue), we propose that more complex tasks (multiple targets) reduce the probability of participants finding and employing a useful strategy. Explicit learning likely draws on working memory as a limited resource (69) and this resource limitation may prevent many participants from storing and combining enough relevant information (color, target direction, force direction/kinematic error) to develop an adequate strategy in complex scenarios. A similar effect of working memory would not be expected to play a crucial role for implicit learning (70, 77). This interpretation aligns with the picture painted by individual participant data: the advantage in learning with a single target appears to arise gradually and across participants with the visual location cue, as would be expected from limited spatial generalization. In contrast, with the color cue, the advantage arises in specific participants and suddenly (e.g. P30), as would be expected from the probabilistic event of finding a strategy. Still, the relative infrequency of “explicit” learners and the limited size of our current sample prohibit strong conclusions on potential effects of task complexity, and this aspect needs further investigation in future studies.

One noteworthy point is the bell-shaped force profiles generated by explicitly adapted participants. This suggests that even participants that used explicit strategies were able to properly compensate for the force-field, similar to the implicit learning participants, even though they were not instructed about the velocity-dependent characteristics of the force field. The question therefore remains whether, with sufficient practice, this explicit adaptation to the force field would be gradually transformed into an implicit adaptation, for example through associative learning.

It is important to highlight the difference between our results using force fields with previous studies using visuomotor rotations. Here we find that the adaptation induced by the visual separation of workspace is primarily implicit. In contrast, visuomotor adaptation in similar conditions was almost entirely explicit (11–13). While it is unclear why this difference exists, we propose a few different hypotheses. One simple possibility is that there is a difference in the adaptation mechanisms between visuomotor rotation and force field adaptation, for example visuomotor rotation elicits strong explicit adaptation with limited implicit adaptation (60), whereas force field adaptation may be primarily implicit. However, it might also be possible that there is a difference in how the visual feedback location is updated in the state representation for visuomotor rotation (78, 79).

Placing our findings within the broader picture of context-dependent learning of dynamics, recent theoretical developments propose a generative model whereby latent contexts give rise to both contextual cues and dynamics. Learners then infer the statistical relations between these and select movements based on prior expectations about context-dependent dynamics after observing cues (80, 81). Comparing this normative framework to the process-based view proposed here is not straightforward and it will be interesting to compare these views in future research. Speculating from our present point of view, context inference in our framework would be expected to show primarily for indirect cues and affect the level that we have operationalized as explicit learning. Inferring the statistical relations with a separate context level would not seem necessary for “implicit” dual-adaptation, as direct cues attribute changes in dynamics to different parts of state space. This aligns with recent findings on generalization (as an inverse of dual-adaptation), which showed that generalization accounts separately for low level kinematic movement features and high level similarity-based context judgment (82). Note that we do not take this to mean that all context inference is explicit - rather, this comparison approaches the limits of our mapping the implicit/explicit dichotomy to a potential hierarchy of motor learning processes.

In summary, we interpret our results as supporting the notion that the sensorimotor system distinguishes different types of cues in context-dependent motor adaptation. One class of cues including visual workspace location enables dual-adaptation directly via state-dependent implicit learning. Another class of cues, including color cues, does not do so, though dual-adaptation with these cues may occur via different routes, such as explicit learning (13, 16, 83). Attempting to categorize cues previously found effective versus ineffective (or inconclusive), the emerging picture is that effective cues like separate locations can be considered directly representative of the dynamic or neural state, thus allocating adaptation to different regions in the state space (40, 84), whereas ineffective cues like arbitrary peripheral movement (31) or temporal sequence (4, 85) are not directly related to dynamic state. Our study thus adds clarification to the role of color cues in this framework by supporting their membership with the latter class and suggesting that previous results that found this cue effective may be explained by explicit learning.

## Notes

### Competing Interest Statement

The authors have declared no competing interest.

### Summary of Updates

This version of the manuscript has been revised to add and update Supplemental Data link and document

